# Effect of the Environmental Mechanical Heterogeneity on T Cell Function

**DOI:** 10.1101/2024.09.28.615572

**Authors:** Jatin Jawhir Pandit, Abed Al-Kader Yassin, Carlos Ureña Martin, Guillaume Le Saux, Angel Porgador, Mark Schvartzman

## Abstract

T cells, key players in the immune system, recognize antigens via T-cell receptors (TCRs) and require additional costimulatory and cytokine signals for full activation. Beyond biochemical signals, T cells also respond to mechanical cues such as tissue stiffness. Traditional *ex-vivo* mechanostimulating platforms, however, present a uniform mechanical environment, unlike the heterogeneous conditions T cells encounter *in-vivo*. This work introduces a novel mechanically heterogeneous environment, with alternating soft and stiff microdomains, to mimic the complex mechanical signals T cells face. Results show that T cells exposed to this heterogeneous environment do not average the mechanical signals but instead respond similarly to those on a homogeneously soft surface, leading to lower activation compared to those on a stiff surface. Interestingly, long-term exposure to these patterns enhances the proliferation of central memory and effector T cell phenotypes, similar to stiff environments. These findings reveal the non-linear nature of T cell mechanosensing and suggest that mechanical heterogeneity plays a critical role in modulating T cell responses, providing new insights into T cell activation and potential implications for immunotherapies.

## Introduction

T cells play a crucial role in the immune system by recognizing and destroying infected or cancerous cells and coordinating the overall immune response. Antigen recognition by T cells is based on T-cell receptors (TCRs), which bind to antigenic peptides presented by major histocompatibility complex (MHC) molecules on the surface of other cells. For T cell activation, this antigen recognition must be accompanied by costimulatory signaling from the interaction between costimulatory molecules on the T cell (such as CD28) and their ligands on the antigen-presenting cells (APCs) or target cells (such as B7), and cytokine signaling produced by the APC or surrounding environment that binds to cytokine receptors on the T cell, providing differentiation and survival signals. Recent evidence indicates that, in addition to these biochemical signals, T cells detect and respond to mechanical cues in their environment, such as the stiffness of the tissue or the mechanical properties of antigen-presenting cells[1–5]. This mechanosensing is crucial for T cell activation and function, as the physical forces and structural properties encountered during interactions with antigen-presenting cells can influence TCR clustering, signal transduction, and ultimately the immune response[6–10]. These studies suggest that mechanosensing helps T cells distinguish between different types of cells[11], ensuring appropriate responses to infections and other immune challenges.

Systematic study of how T cells respond to mechanical cues can be based on their ex vivo activation in mechanostimulating environments with varied elasticity, commonly using elastomeric gels functionalized with antigen molecules[12,13]. Various reports have shown different types of dependence of T cell response to variations in elasticity, such as either an increase[14,15] or decrease in T cell activation with increased elasticity[16]. The trend in T cell activation changes with environmental elasticity highly depends on the elasticity range and the experimental conditions under which T cells are stimulated. It is clear that the studies of T cell response to environmental elasticity so far only scratch the surface of the complex mechanisms of T cell mechanosensing and mechanotransduction, which are yet to be fully elucidated.

One limitation of the state-of-the-art elastomer-based mechanostimulating platforms for T cells is that they provide a homogeneous mechanical environment, giving T cells a uniform type of mechanical signaling. This contrasts with the physiological environment of T cells, characterized by high mechanical heterogeneity. In vivo, T cells interact with relatively soft lymphoid organs like lymph nodes and the spleen, as well as peripheral tissues such as skin and muscles with various degrees of stiffness. Also, APCs such as dendritic cells, macrophages, and B cells, and endothelial cells within blood vessels provide T cells with different mechanical stimuli, often simultaneously. A fundamental question arises: when a T cell is exposed simultaneously to multifaceted mechanical environments and receives different mechanical signals, does it average these signals and produce an average response, or does one signal dominate? Addressing such questions requires an artificial mechanostimulating environment that simultaneously provides different mechanical signals. However, such engineered microenvironment has not yet been demonstrated.

In this work, we studied the effect of environmental mechanical heterogeneity on T cell immune response. To that end, we engineered a model environment for mechanical heterogeneity consisting of surfaces patterned with alternating soft and stiff microdomains, sized below the size of T cells (Fig. 1a). This design ensured that a single T cell, when in contact with such a surface, would necessarily be exposed to mechanical signals from both microdomains. We activated primary human T cells on these mechanical patterns and compared their response to that of T cells activated on homogeneously soft and stiff surfaces. We found that T cells activated on the stiffness patterns do not average the mechanical signals produced by the microdomains with varying stiffness, but rather produce relatively low activation equal to that of the homogeneous soft surface, while the homogeneous stiff surface caused the highest activation. We then studied the effect of mechanical heterogeneity on early signaling but found it to be similar to that produced by homogeneous mechanical surfaces. Finally, we tested the long-term effect of environmental mechanical heterogeneity on T cell function by characterizing T cell proliferation and differentiation. In particular, we found that the stiffness patterns enhance the proliferation of central memory and effector phenotypes of T cells, as wells as the stiff surfaces do, as compared to soft surfaces. Overall, these results are the first of their kind to demonstrate the non-linear response of T cells to multiplexing of mechanical signals produced by a heterogeneous environment, providing important insight into the mechanical sensing of T cells.

**Figure 1.**
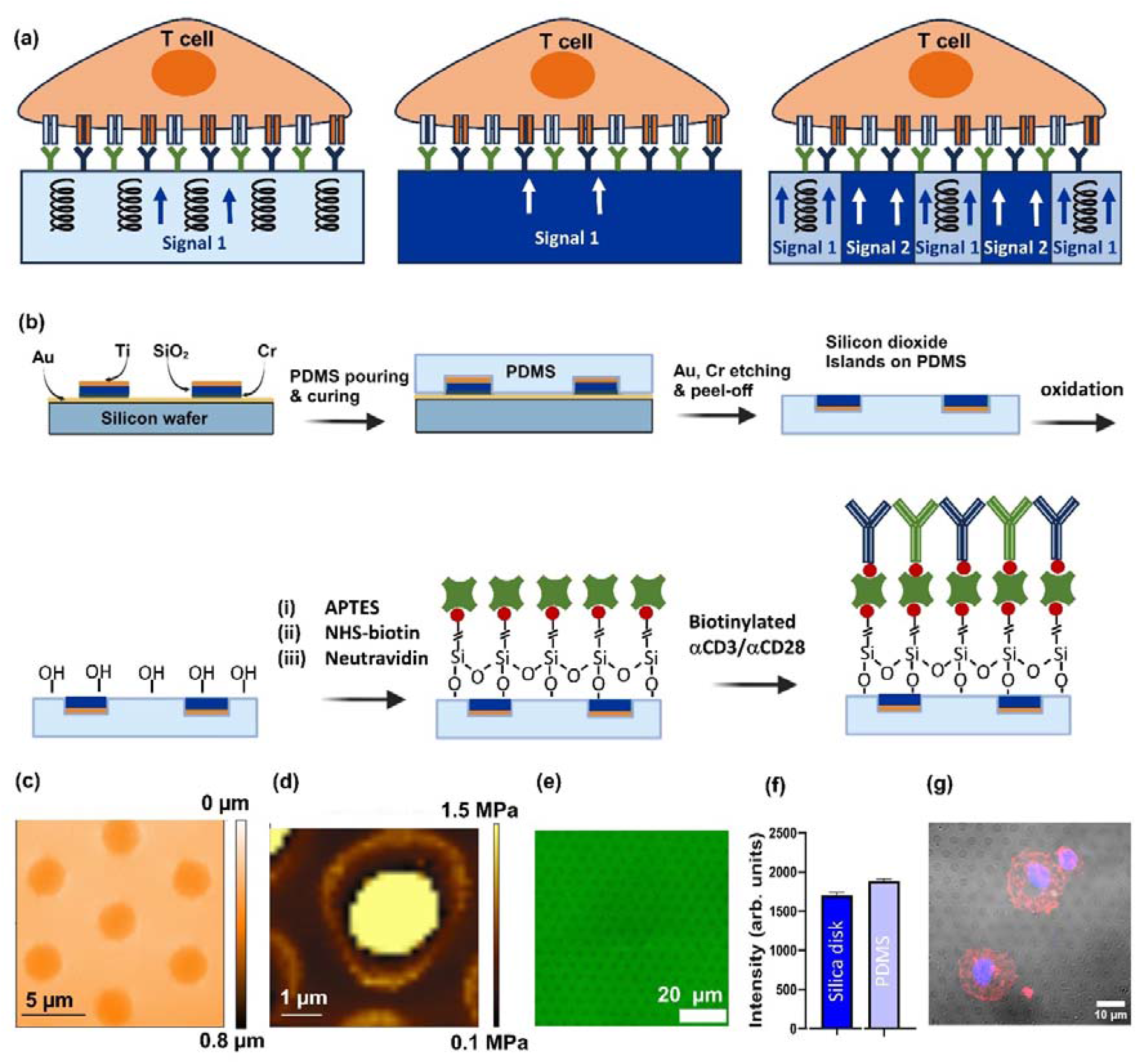
(a) Scheme of two activating microenvironment – soft and stiff, each providing uniform mechanical activating signal, vs. mechanically patterned microenvironment simultaneously providing different mechanical signals to each cell. (b) Fabrication and biofunctionalization process flow of the stiffness pattern. (c) Topography AFM scanning of the stiffness pattern. (d) Modulus AFM mapping of the stiffness pattern. (e) Fluorescence microscopy image of the stiffness pattern after the biofunctionalization, showing the presence of tagged Neutravidin on both PDMS and Silica domains. (f) Quantitative comparison between the fluorescence signals from PDMS and Silica domains. (g) Primary Human T cells spread on fluorescence pattern. The cells were incubated on the pattern for 24 hrs, fixed and stained for cytoskeleton with phalloidin (red) and for nucleus with DAPI (blue).

## Results

Stiffness patterns were based of hexagonal array of SiO_2_ discs surrounded by soft Polymethyl Siloxane (PDMS). The discs were 3 μm in diameter, and were arranged with the periodicity of 6 μm . The arrays were fabricated by producing stiff discs on sacrificial surface based on Silicon covered with Au thin film, by standard photolithography, electron beam evaporation of Cr/ SiO_2_/Ti, and liftoff (Fig. 1 b). The disks were then mechanically transferred to PDMS, by PDMS pouring and curing, followed by mechanical detachment from Silicon, and wet etching of Au and Cr (details appears in methods section). Since T cells are around 10 μm in diameter, this pattern geometry ensures that each cell approaching the surface is exposed to both the soft and stiff domains. Following the fabrication, the entire surface – including Silicon Dioxide and PDMS domains - were uniformly functionalized to provide T cells with the essential activating and costimulatory biochemical stimuli. To that end, the surface was first treated with Oxygen plasma, and following by chemisorption of (3-Aminopropyl) trietoxysilane (APTES), and subsequent attachment of biotinyl succinimide, to which even mixture of biotinylated antibodies against CD3 and CD28 receptors were attached via NeutArvidin bridge .

The fabricated surfaces were characterized by AFM (Fig. 1 c), prior the biofunctionalization. A slight topography of ∼140 nm (Fig. S1) was formed by Silicon oxide discs, most probably due to the shrinkage of PDMS during its curing. Notably, high-aspect ratio topographic features with a few hundred nm diameter and micron scale depth were recently shown to induce T cell activation without any biochemical stimulation[17]. On the other hand, the topography obtained here was negligible, and by itself cannot produce any effect of T cells. Thus, in the context of T cell activation, the fabricated surfaces can be considered flat. Also, force-distance AFM scanning was used to characterize the stiffness heterogeneity of the obtained surface pattern (Fig. 1d). The measured elastic modulus of PDMS was ∼100 Kpa, while the modulus obtain within the silicon dioxide disc was ∼1.5 MPa. Of course, the latter result does not represent the bulk modulus of Silicon dioxide, which is typically within the range of tens GPa. This result can be explained by the fact that the deflection of the surface within the relatively thin (∼30 nm) Silicon Oxide disc upon the force applied by AFM tip is greatly affected by the undelaying soft PDMS. Still, the obtained elasticity map clearly demonstrates the existence of a high contrast between the soft (PDMS) and stiff (Silicon dioxide) domains in terms of their response to mechanical force. After the biofunctionalization, the uniformity of the molecular coverage was confirmed by fluorescence microscopy, by imaging NeutrAvidin™ that was fluorescently tagged with Oregon Green™ 488 (Fig. 1e). The fluorescent intensity measured on both Silicon Dioxide and PDMS domains indicate that both were covered with densely packed monolayers of NeutrAvidin, ensuring uniformly high biochemical stimuli on these domains provided by activating and costimulatory antibodies (Fig. 1 f).

After the confirmation of the stiffness heterogeneity and the biofunctionalization uniformity, the patterned surfaces were used to study the response of T cell to the environmental mechanical heterogeneity. To that end, human Peripheral Blood Mononuclear Cells (PBMCs) freshly isolated from three healthy donors were seeded onto these surfaces and were incubated for 4 hrs. Also, the cells were seeded on two control surfaces : (i) homogeneous soft control made of PDMS with the same composition and properties as that used in the stiffness pattern, and (ii) stiff control based of glass cover slip. Both controls were functionalized with a-CD3 and a-CD28, similar to the stiffness pattern, to deliver the same biochemical signals for T cell activation. Fig. 1 g shows the confocal images of cells on the patterned surfaces; the cytoskeleton of the cells (red) and the nucleus (blue) were visualized. These images revealed that the cells adopted the typical morphology of adherent lymphocytes developing protrusions; these protrusions were seen on soft as well as stiff parts of the sample, confirming interaction with the patterned surfaces[18].

To study whether and how the stiffness heterogeneity of the activating surface affects the response of T cells, we first assessed their expression of Lysosomal-associated membrane protein-I (CD107a). T cell activation is followed by degranulation, in which lytic granules containing perforin and granzyme[19] diffuse to the membrane, release their lytic content, bringing CD107a to the surface[20]. This makes CD107a a commonly used marker for the activation of cytotoxic T cells. Here, PBMCs were seeded on the stiffness patterns and the control surfaces for four hours, and stained for CD4, CD8, and CD107a, and analyzed by flow cytometry. Plastic surface without stimulating antibodies was used here as negative control. Fig. 2 a and b show typical degree of CD107a expression, separately for CD4+ and CD8+ subsets, and Fig. 2 c and d show the % of CD107a positive cells across the donors. Notably, it has been generally considered that the role of CD8+ T cells is to directly eliminate pathogens, and of CD4+ T cells – to contribute to the immunity by helping and CD8+ T cells, B cells, Natural Killer cells. However, it is evident nowadays that CD4+ T cells can, to certain extent, perform cytotoxic functions and cause direct apoptosis of target cells[21]. Still, the overall expression of CD107a by CD4+ T cells, as anticipated, is substantially lower than that by CD8+ T cells. Nevertheless, both subsets demonstrate the same pattens of CD107a expression consistent for all the donors, by which stiff control samples produced the highest degranulation. In contrast to stiff surfaces, both stiffness pattern and soft control produced similarly lower levels of degranulation, for both CD4+ T cells and CD8+ T cells, consistently for all the tested donors.

**Figure 2:**
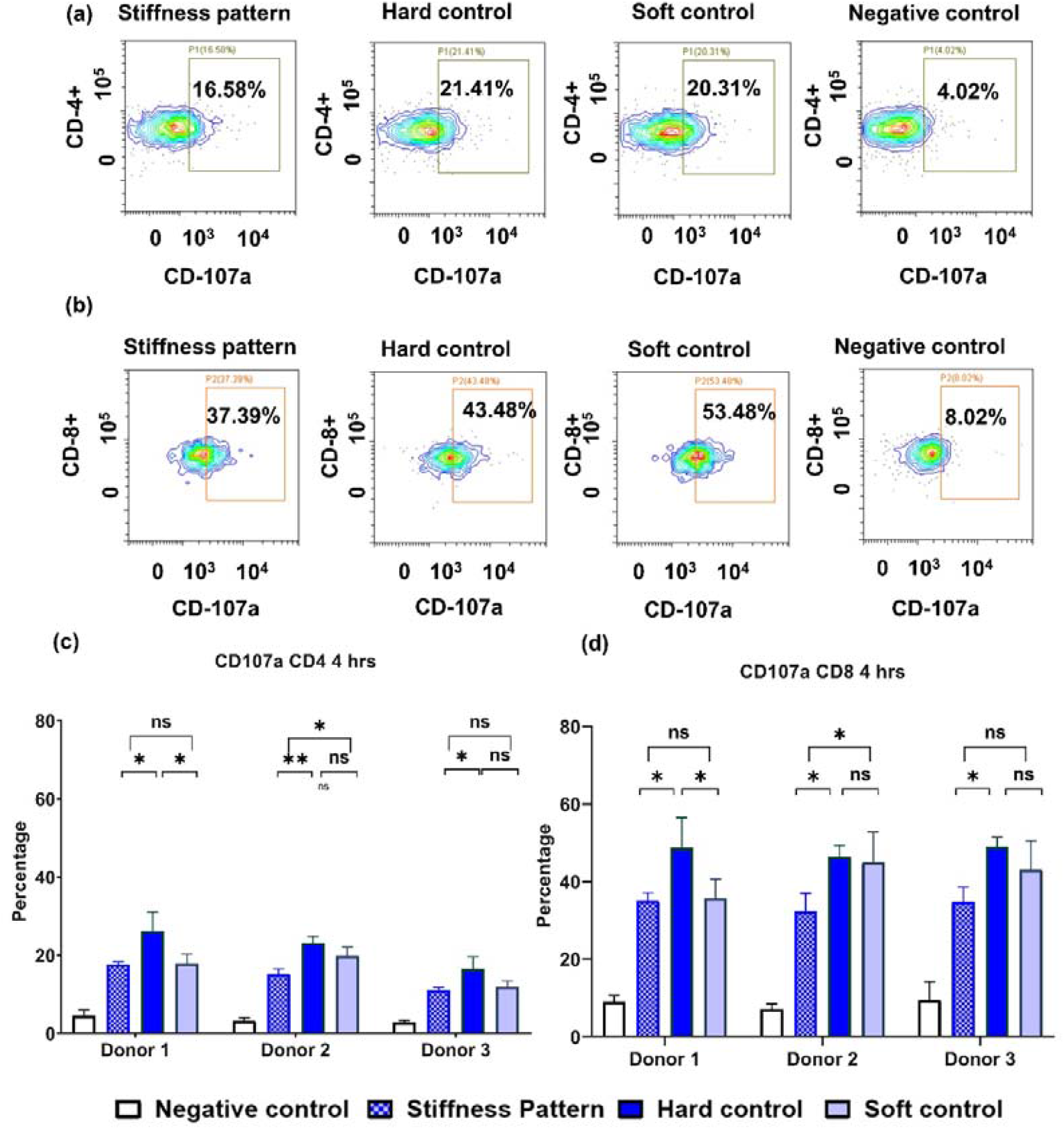
CD107a expression after 4 hrs incubation. Dot plots of the flow cytometry are shown in (a,b) Graphs (c,d) show the expression of CD107a for CD4^+^ and CD8^+^ cells, respectively. The statistical analysis was performed in Tukey’s multiple comparison test using GraphPad Prism software. *p < 0.05, **p < 0.01, ***p < 0.001, ****p < 0.0001, and ns: not significant

We next tested whether and how the type of the activating surface affects another activation marker -- CD69. CD69 is a membrane-bound, type-II lectin receptor that rapidly appears on the plasma membrane after TCR/CD3 engagement, and hence serves as an early activation marker8. Herre, PBMCs were incubated for 24 hrs, stained against CD69, CD4, and CD8, and analysed using flow cytometry (Fig. 3) . The graphs shown in Fig 3 (c,d) for CD69 taken after 24 hrs corroborated with the findings from CD107a after 4 hrs for two of the three donors, with the stiff control having the highest activation and the patterned sample and the soft control having little to no significant variation in activation. Stil, even for these two donors, the increased CD69 expression for stiff surfaces was not always statistically significant. Overall, the effect of stiff surface on the expression CD69 was less pronounced than in the case of CD107a. On the other hand, stiffness pattern produced the same CD69 level as the soft control, mirroring the results of by CD107a expression.

**Figure 3:**
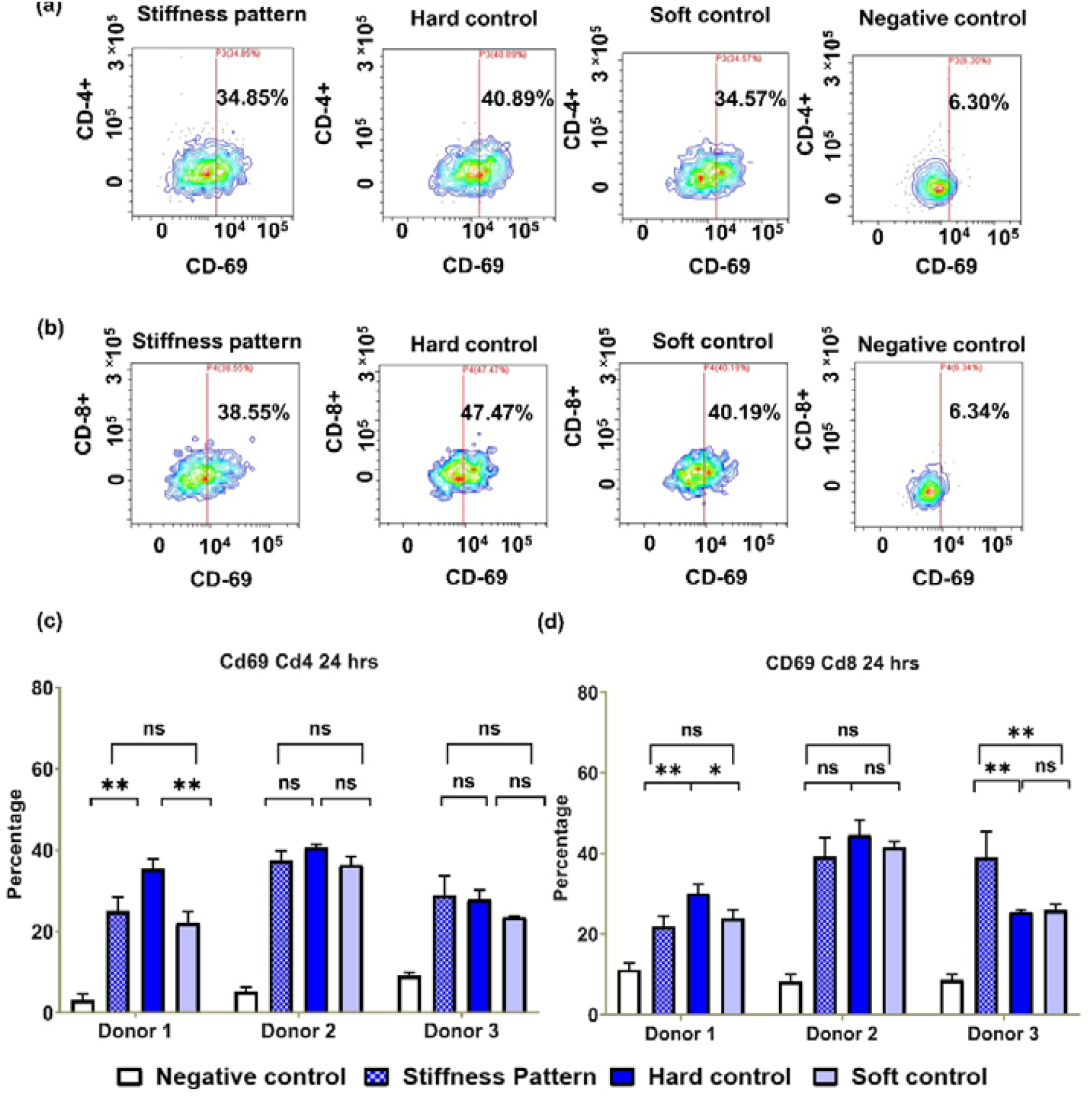
CD69 expression by T cells three donors, after 24 hours incubation. Dot plots of the flow cytometry are shown in (a,b), the Graphs (c,d) show the expression of CD69 for CD4+ and CD8+ cells, respectively. The statistical analysis was performed in Tukey’s multiple comparison test using GraphPad Prism software. *p < 0.05, **p < 0.01, ***p < 0.001, ****p < 0.0001, and ns: not significant.

Next, we investigated how stiffness heterogeneity affects early signaling in T cells. Activation of the TCR signaling cascade is associated with the binding of the immune receptor activation motif (ITAM) by ZAP-70, which is further phosphorylated by Lck. This phosphorylation activates ZAP-70 catalytic activity and its autophosphorylation, initiating downstream signaling and serving as an early signaling marker. To assess the intensity of ZAP-70 phosphorylation, PBMCs were seeded onto the surfaces for 15 minutes, fixed, and stained for ZAP-70 phosphatase using indirect immunofluorescence staining. The cells were imaged using a fluorescence microscope, and the phosphorylation intensity was quantified by fluorescence signal. Fig. S2 (a, b, c) shows microscope images of T cells on stiff control, soft control (PDMS), and stiffness patterns, stained for the cytoskeleton, nuclei, and phospho-ZAP-70. The graph in Fig. S2 (d) shows the fluorescence intensity of ZAP-70 phosphorylation on different surfaces. The results indicate that the stiff control had slightly higher phospho-ZAP70 intensity than the soft control and the pattern, although the difference is not statistically significant. Additionally, the location of phosphorylated Zap-70 molecules within cells activated on the stiffness pattern does not specifically correlate with either the soft or stiff domains, indicating that the stiffness distribution of the activating surface has no observable effect on early signaling.

In contrast to the negligible effect of the activating surface on ZAP-70, the effect of the activating surface on the T cell morphology is unignorable. Fig. 4a show T cells spread of the three used surfaces after 15 minutes of stimulation, which show typical isotropic shape, indicating string interaction with the surface in each case. The distribution of acting is peripheral , which is typical for well spread and adhered cells, yet this peripherality is mostly pronounces for stiff surfaces. On the patterned surfaces, the actin was distributed in more random fashion, showing little sub-micron areas with high actin concentrations. However, there were no clear correlation between the areas of actin concentration and the soft or stiff domains of the patterned surface underneath the cells. We also characterized the cell area, which is one of the main merits of the cell morphology, and is an indicator that the T cell receives sufficient signals to become fully activated and effectively respond to the presence of antigens. T cell spreading as the function of the elasticity of the undelaying surface was previously studied, and the observed values of the area, as well as trends of the cell area vs. elasticity were highly dependent on the cell type (e.g. primary T cells vs. T cell line), range of probed elasticities, and other experimental conditions such as presence of activating, costimulatory, and adhesive molecules on the surface[6,15,22–25]. Here, we found that the stiffness pattern produced an average cell area slightly higher that the stiff controls did, yet without statistically significance (Fog. 4b). On the other hand, both the stiffness pattern and stiff controls produced higher cell area than the soft control. The type of the activating surface also influenced the circularity of the cells: here, as anticipated, the stiff surfaces produced the most circular cell shapes, on average, that the stiffness pattern and soft control (Fig. 4c).

**Figure 4.**
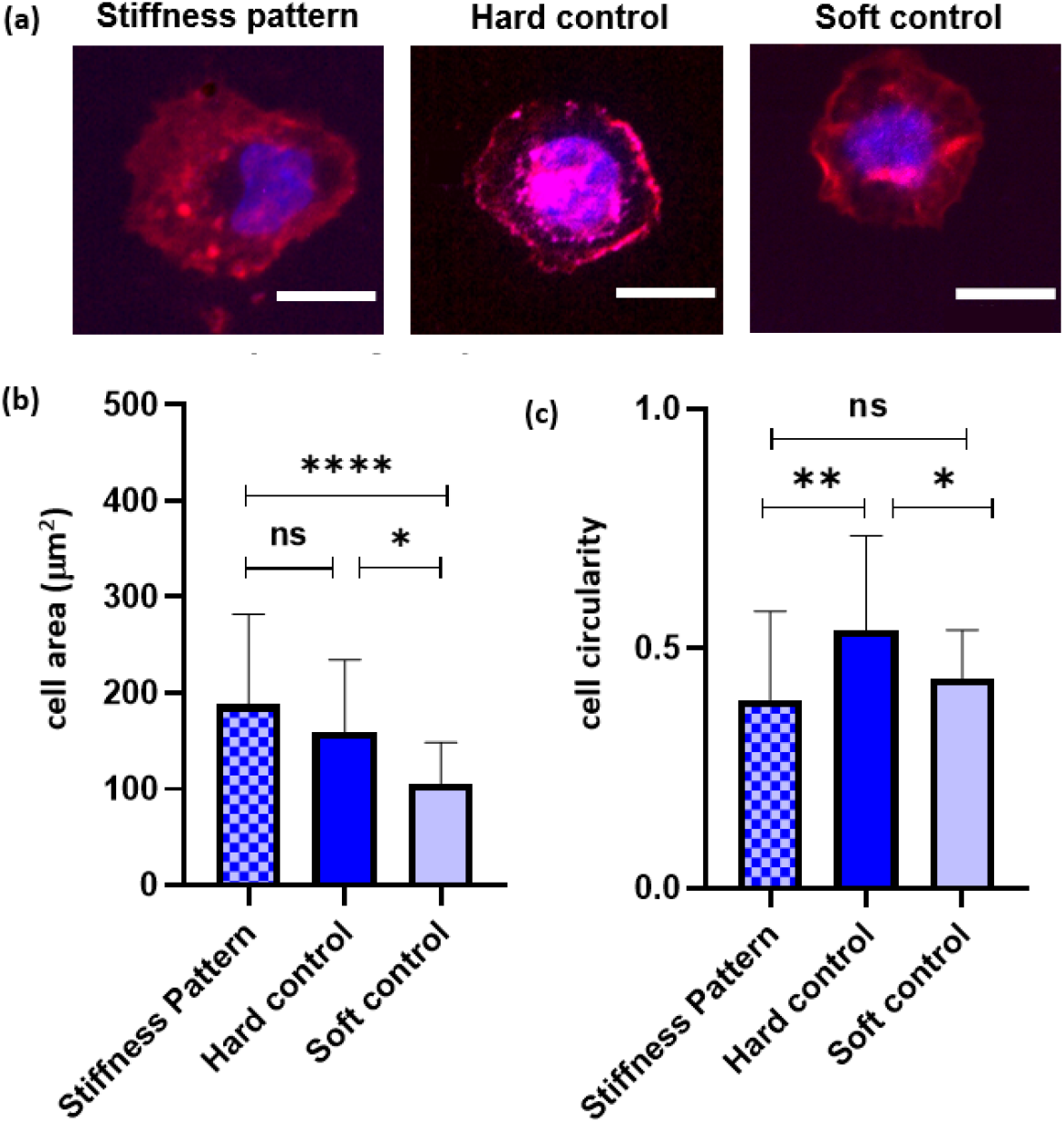
(a) Typical T cells spread on different probed surfaces. The cells were spread for 15 minutes, then fixed and stained for cytoskeleton with phalloidin (red) and DAPI for nucleus (blue). (b) Average cell area on the probed surfaces. (c) Average cell circularity on the probed surfaces. The images were taken by confocal microscope, and analyzed using ImageJ software. Between 30 to 40 cells were analyzed for each type of surfaces. The statistical analysis was performed in Tukey’s multiple comparison test using GraphPad Prism software. *p < 0.05, **p < 0.01, ***p < 0.001, ****p < 0.0001, and ns: not significant

After investigating the effect of mechanical heterogeneity of the stimulating environment on T cell activation and morphology, we tested whether this heterogeneity affects T cell differentiation. For this purpose, we stimulated the cells on the tested surfaces for one day, then transferred them into new wells, and incubated for four days. On Day 4 and Day 7, the proliferated cells were T cells stained for CCR7 and CD45RO, and their expression was analyzed by flow cytometry. These markers are commonly used to differentiate between different phenotypes of T cells. Naïve T cells express CCR7, which helps them in their migration to lymphoid organs in search of antigens presented on APCs[26]. CD45 is a transmembrane tyrosine phosphatase found on all cells of the hematopoietic lineage except red blood cells (RBCs); memory T-cells express a unique isoform of CD45 known as CD45RO[27]. Both these markers are used in combination to analyze various T cell subsets, and, while analyzed together, divide the T-cells into four different subsets: naïve subset (CCR7^+^CD45RO^−^ cells), central memory (CCR7^+^CD45RO^+^ cells), effector memory (CCR7^−^CD45RO^+^ cells), and terminal differentiated effector memory (TEMRA, CCR7^−^CD4RO^−^ cells)[28].

The results in Figure 5 show the fold expansion of T cells in general (based on counting CD3+ T cells), and for different T cell phenotypes for the three tested donors. First, it can be concluded that the type of surface insignificantly affected the general T cell proliferation (Fig. 5 a-b). In all cases, a 15 to 25-fold expansion was achieved after 7 days, with no clear dependence on the activating conditions. Looking separately at central memory T cells (Fig. 5 c-f), there is an effect of the activating surface: on day 4, both patterned and stiff surfaces resulted in higher differential proliferation of CD4^+^/CD8^+^ T_cm_ as compared to the soft surface, for all the three donors. Similarly, on day 4, both patterned and stiff surfaces resulted in higher differential proliferation of CD4^+^/CD8^+^ T_em_ as compared to the soft surface. Two other phenotypes – TEMRA and Naïve, demonstrated substantially lower proliferation, as anticipated (Fig. S3). Note that the differences for T_em_ and T_cm_, between patterned and stiff surfaces were either insignificant or relatively small. Therefore, we can conclude that the presence of stiff surface, either in a patterned/non-patterned form, accelerate the proliferation of these two T subsets as compared to naïve T cells. Yet, following additional days of culture, the fraction of these two subsets reaches saturation and no difference between the activation surfaces.

**Figure 5.**
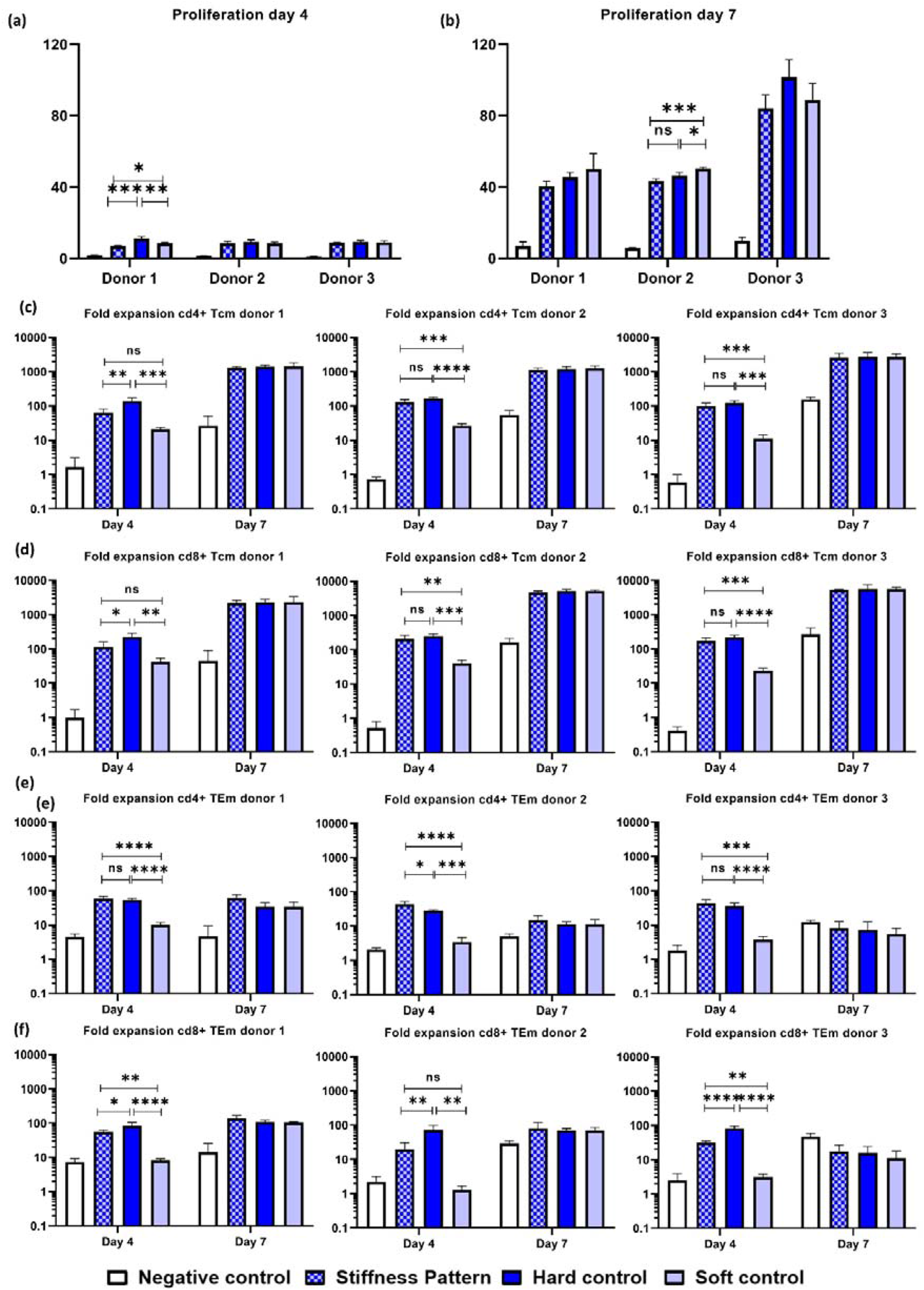
(a), (b) Overall proliferation of T cells on day 4 and day 7, respectively, vs. the type of the activating surface. (c) Proliferation of CD4+ T_cm_ Phenotype, (d) Proliferation of CD8+ T_cm_ Phenotype, (e) Proliferation of CD4+ T_em_ Phenotype. (f) Proliferation of CD8+ T_em_ Phenotype. The statistical analysis was performed in Tukey’s multiple comparison test using GraphPad Prism software. *p < 0.05, **p < 0.01, ***p < 0.001, ****p < 0.0001, and ns: not significant .

## Discussion

The mechanically heterogeneous environment of T cells encompasses a wide range of physical properties and forces that T cells encounter as they navigate through different tissues and interact with various cells in the body. This heterogeneity significantly influences T cell behavior, activation, migration, and overall immune response. Lymphoid organs like lymph nodes and the spleen provide a relatively soft and elastic environment that facilitates T cell activation and interaction with other immune cells[29]. This softer environment supports the rapid movement and scanning of T cells for antigens presented by dendritic cells and other APCs[30]. In contrast, peripheral tissues such as skin, muscles, and connective tissues exhibit varying degrees of stiffness. For example, the extracellular matrix (ECM) in these tissues is often denser and more rigid, requiring T cells to exert greater force to migrate and infiltrate these areas[31,32]. The mechanical properties of these tissues can affect the efficiency of T cell movement and their ability to reach infection or tumor sites.

The mechanical properties of cells presenting antigens for T cell activation, including professional APCs and target cells, play a crucial role in the formation of the immunological synapse and the effectiveness of antigen presentation. T cells can exert mechanical forces to probe the stiffness of encountered cells, influencing T cell receptor (TCR) signaling and activation[7]. The mechanism of TCR activation is still greatly debated, yet in all proposed models, the effect of force on TCR triggering is critical. In the allosteric-conformation-based model, TCR binding to a ligand leads to the release of the CD3 cytoplasmic tail from the inner membrane leaflet, resulting in phosphorylation of the immunoreceptor tyrosine-based activation motif[33,34]. The details of how extracellular ligand binding causes changes in intracellular conformation are still unclear, but it has been proposed that the force applied to the receptor changes the conformation of the transmembrane domain of the TCR α-subunit and disrupts the association with CD3ζζ cytoplasmic chains, resulting in increased accessibility for ζζ phosphorylation[35,36]. Alternatively, the kinetic segregation model does not consider TCR-CD3 conformational changes but explains TCR triggering by the physical segregation of large phosphatase molecules, such as CD45, from the binding domain. This segregation shifts the kinase-phosphatase balance towards kinase activity, initiating activating signaling cascades[37–39]. Due to the catch bond nature of TCR-ligand binding[40], where mechanical force facilitates bond affinity, force applied in this context stabilizes the physical proximity between the two membranes at the binding region, thereby stabilizing CD45 exclusion and enhancing the generation of activating signaling.

This work provides two important insights into how the mechanical environment regulates T cell responses. First, it shows that multiplexing mechanical signaling by exposing T cells to mechanically heterogeneous environments produce a non-linear response. Here, the stiff control surface produced a higher activation stimulus than the soft surface, as reflected by both CD69 and CD107a markers. However, a heterogeneous stiffness pattern that a priori provided cells with both types of stimuli produced activation similar to that of the soft surface. The reason for this behavior is yet to be elucidated. One possible explanation is based on the finding that stiffer substrates produce higher deformation of the T cell membrane and thus better exclusion of CD45 from the binding regions, leading to a greater number of TCR-ligand interactions[15]. The stiffness pattern design aimed to produce both types of mechanical stimuli from soft and stiff domains. Yet, the soft domain appeared in this pattern as a continuum, and thus could dominate in determining TCR-CD45 segregation across the long physical range of the T cell-surface interface. On the other hand, stiff domains are discontinuous and thus control TCR-CD45 segregation within the physical range limited by the domain size, limiting their influence on the overall T cell response. Verifying this explanation requires further research, which could include microscopic visualization of CD45-TCR segregation at the T cell interface on soft and stiff domains. Additionally, studying T cell responses to the inverse stiffness pattern, with soft microdomains surrounded by a stiff continuum, could shed light on the mechanism of heterogeneous mechanical stimulation.

Previous studies were demonstrated that the stiffness of activating surface determines the proliferation of T cells, with the general tendency of the proliferation increase with the decrease in the stiffness[16,41]. Yet, the observed trend of stiffness vs. proliferation largely depends on the proliferation conditions, and on the probed stiffness range Note that the response of T cells on synthetic surfaces is biphasic: it increases with the stiffness up to a peak of a few tens KPa (which corresponds to the stiffness of antigen presenting cells), and then decreases[6,22]. In contrast, PDMS elasticity used in this research was about 100 kPa – which is above the optimal elasticity for T cell activation, and this can explain relatively lower proliferation it produced on day four. This work, and specifically the new fabrication methodology of stiffness patterns presented here, paves the way for follow-up studies on the effect of heterogeneous mechanical environment structures of elements of different stiffnesses beyond those explored here, including but not limited to antigen presenting cells.

Besides the important insights this work provides on mechanosensing, it introduces significant novelty in the realm of micro/nanostructured materials for ex vivo T cell stimulation. These materials have been used so far to understand how ligand distribution affects T cell spreading, adhesion, and immune response [6,42–45]. Furthermore, micro/nano-structured materials have been employed to study how the physical features of the cellular environment, with a focus on elasticity and microtopography, regulate the activation and function of T cells[9,12,46–48] as well as other lymphocytes, such as NK cells[44,47,49–51]. An emerging area in this field involves novel material based approaches for immunotherapy, including, but not limited to, nanomaterials for priming immunotherapeutic T cells and scaffolds that provide a stimulatory microenvironment for adoptive cell transfer to expand T cells and maintain their antitumor activity[52–54] . This work makes an additional important contribution to the study of engineered materials for T cells by introducing the possibility to simultaneously deliver mechanical signals to T cells and investigate the effects of signal multiplexing on T cell response. Moreover, the proposed new fabrication approach is not limited to T cells and will undoubtedly pave the way for numerous follow-up studies aimed at understanding the effects of mechanical diversity on different cells and biological systems.

## Materials and methods

### Fabrication of Heterogenous stiffness surface

Silicon wafers were cut into 2.5 by 2.5 cm squares and were cleaned by sonicating in acetone for 5 minutes, followed by rinsing with ethanol. Cleaned samples were then sputtered with 20nm of gold before proceeding with the photolithographic process. The samples were then spin-coated with AZ1505 positive photoresist (Microchemicals GmbH) and a hexagonal array of disks having a diameter of 3 microns separated by 3 microns edge-to -edge was produced. Samples were then moved to E-gun evaporator for deposition of Cr (4nm), SiO2 (30nm), and Ti (3nm), respectively. After lift-off, by sonication in acetone, the samples were then mounted on Sylgard 184 PDMS with stiffener to base ratio of 1:50. The mounted samples were degassed and then left to cure in the oven overnight at 60°C. After removal of gold and chromium using appropriate etchants (both from Merck), the PDMS was peeled the silicon wafer to produce the patterned elastomeric substrates.

#### Biofunctionalization of the heterogeneous stiffness patterns

The surfaces, after being activated by UV ozone for 5 minutes, underwent treatment with 3-aminopropyl)-triethoxysilane (APTES, Sigma-Aldrich) by immersing them into 5% ethanolic APTES solution for 30 minutes at room temperature and then washing with ethanol before baking in an oven at 60°C for 30 minutes. After APTES modification, surfaces were further functionalized with biotin by immersion in a 1 mM aqueous solution of biotinyl succinimide ester (EZ-Link NHS-Biotin; Thermofisher) overnight. Subsequent steps were performed in a sterile hood using sterile buffers.

After rinsing the samples with Phosphate Buffer Saline (PBS 1X), the samples were incubated with NeutrAvidin™, Oregon Green™ 488 conjugate (Invitrogen) for 90 min at room temperature. The samples were rinsed with PBS Tween 20 (PBS-T), and then a mix of activating ligands Biotin anti-human α-CD3 and α-CD28 (BioLegend) at a concentration of 2μl/ml in 1X PBS in a ratio of 1:1 was added to the samples and incubated at room temperature. The samples were then rinsed with sterile PBS before cell seeding.

#### AFM Microscopy

Atomic Force Microscopy (AFM) images were acquired using Nanosurf AFM (Dynamic mode, Multi75AI-G tip), and Young’s modulus on different locations on the samples was quantified using ANA software using the Pyramidal Regular 4-sided model. AFM force mapping (Static mode, Multi75AI-G tip) was also performed on the samples, generating a force map to visualize the contrast in stiffness across the pattern.

#### Cell Isolation and culture

Peripheral blood mononuclear cells (PBMCs) were isolated from fresh blood from 3 healthy adult donors recruited by written informed consent, as approved by the Institutional Review Board of the Ben-Gurion University of the Negev. PBMCs were isolated from fresh blood by Ficol gradient, and seeded seeded onto the biofunctionalized samples in 4Cell Nutri-T media (Sartorius) containing <2% serum and 50 units of IL-2 and left to incubate overnight. Silicon dioxide (SiO2) and soft PDMS (stiffener to base ratio of 1:50) were used as controls; in addition to this, internal control was taken, having cells without any activation.

#### Flow Cytometry

For Flow cytometry measurements, 30,000 cells per well were used. The cells were washed with 1X PAF (PBS-0.05% Sodium Azide-2% FCS) and seeded in 96 well plates. The cells were then stained with PE anti-human CD3 Antibody, FITC anti-human CD4 Antibody, APC/Fire™ 750 anti-human CD8 Antibody, PerCP anti-human CD69 Antibody, and APC anti-human CD107a (LAMP-1) Antibody, all acquired from BioLegend at a concentration of 1μg/ml and incubated on ice for 30 minutes. Thereafter, the cells were centrifuged, and the supernatant was discarded before adding DAPI solution (1μg/ml) and incubated on ice till the reading was taken. All the samples were analyzed in Beckman CytoFLEX LX flow cytometer. For overall proliferation analysis, the fraction of CD3-positive cells was calculated, and CD3-positive cells were then analyzed for staining with the other antibodies employed for staining.

#### Immunostaining for Phospho-ZAP-70

Freshly isolated PBMCs were seeded onto the samples and incubated for 15 minutes at 37°C. The samples were then gently rinsed with PBS once and fixed with 4% paraformaldehyde for 15 minutes at 4°C. The cells were then permeabilized using 0.1% Triton-X100 in PBS for 3 minutes at 4°C before moving the samples into ice-cold methanol for 10 minutes at -20°C. The sample was blocked using 2% Bovine Serum Albumin (BSA) in PBS for one hour and then incubated with phosphor-ZAP-70 (1:50 in 2% BSA in PBS) overnight at 4°C. After that, the samples were rinsed with PBS thrice and incubated with Alexa Flour 647 (1:40) and Alexa Flour 555 (1:1000) (Life Technologies) overnight at 4°C. Before mounting the samples with ProLong Gold antifade reagent containing DAPI (Life Technologies), the samples were rinsed with PBS twice and once with deionized water.

### Confocal Microscopy

Freshly isolated PBMCs were seeded onto the fabricated samples and were incubated for 4 hours at 37°c. The incubated samples were then fixed with 4% Paraformaldehyde for 15 min on ice. The fixed samples were then gently rinsed and kept in a blocking buffer (5% BSA in 1X PBS) for 1 hour at room temperature. After blocking, the samples were incubated in antibody coating buffer (1% BSA, 0.1% Saponin, Alexa Flour Phalloidin 555 (1:1000) (Life Technologies) in 1X PBS for 1 hour at room temperature in the dark. After incubation, the samples were rinsed with PBS 2-3 times and mounted with ProLong Gold antifade reagent containing DAPI (Life Technologies). The samples were then visualized using a Zeiss LSM 880 Confocal Laser Scanning Microscope. Image analysis was done using ImageJ software.

### Statistics

Fluorescence intensity measurements were done by taking three different regions on different surfaces with the same exposure time and magnification. All biological experiments were performed three times. 20 to 40 cells were imaged and averaged for quantification. Statistical analysis was performed by analysis of variance, and Tukey’s multiple comparison post hoc test was also performed using GraphPad Prism software. The results were considered to be significantly different for P < 0.05.

## Acknowledgements

This work was supported by German Research Foundation (Project SM 289/10-1. 488) and Israel Science Foundation 489 (Project 2016/21).

