## Supporting Information FIle for "Effect of the Environmental Mechanical Heterogeneity on T Cell Function"

**Effect of the Environmental Mechanical Heterogeneity on T cell Immune Response**

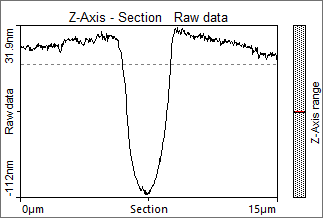

Fig. S1 . AFM Scanning profile of the pattern showing a topography across the Silica disk embedded in PDMS surface

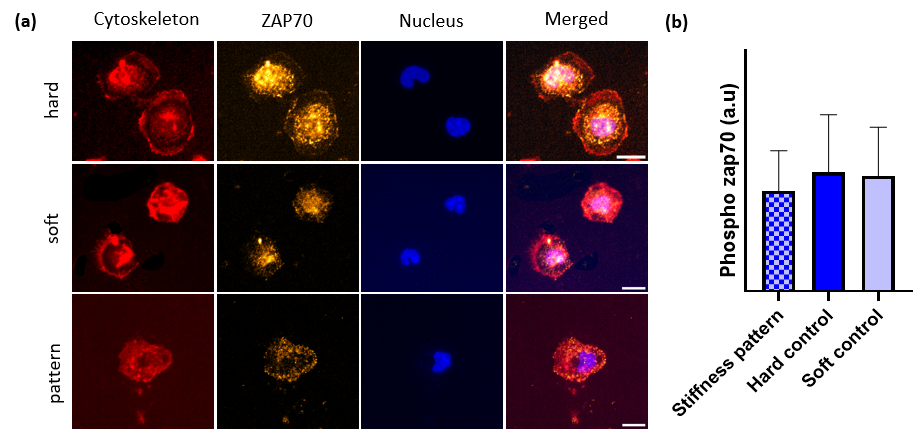

Fig. S2. Early signaling of T cells on various activating surfaces: (a) typical images f T cells 15 minutes afters the exposure to the surfaces. The cells wee fixed and stained for cytoskeleton with phalloidin (red) , for nucleus with DAPI (blue), and for phospho-ZAP70 with tagged anti- phospho-ZAP70. (b) average expression of phospho-ZAP70 for T cells activated on different surfaces, showing no significant effect of the surface type.

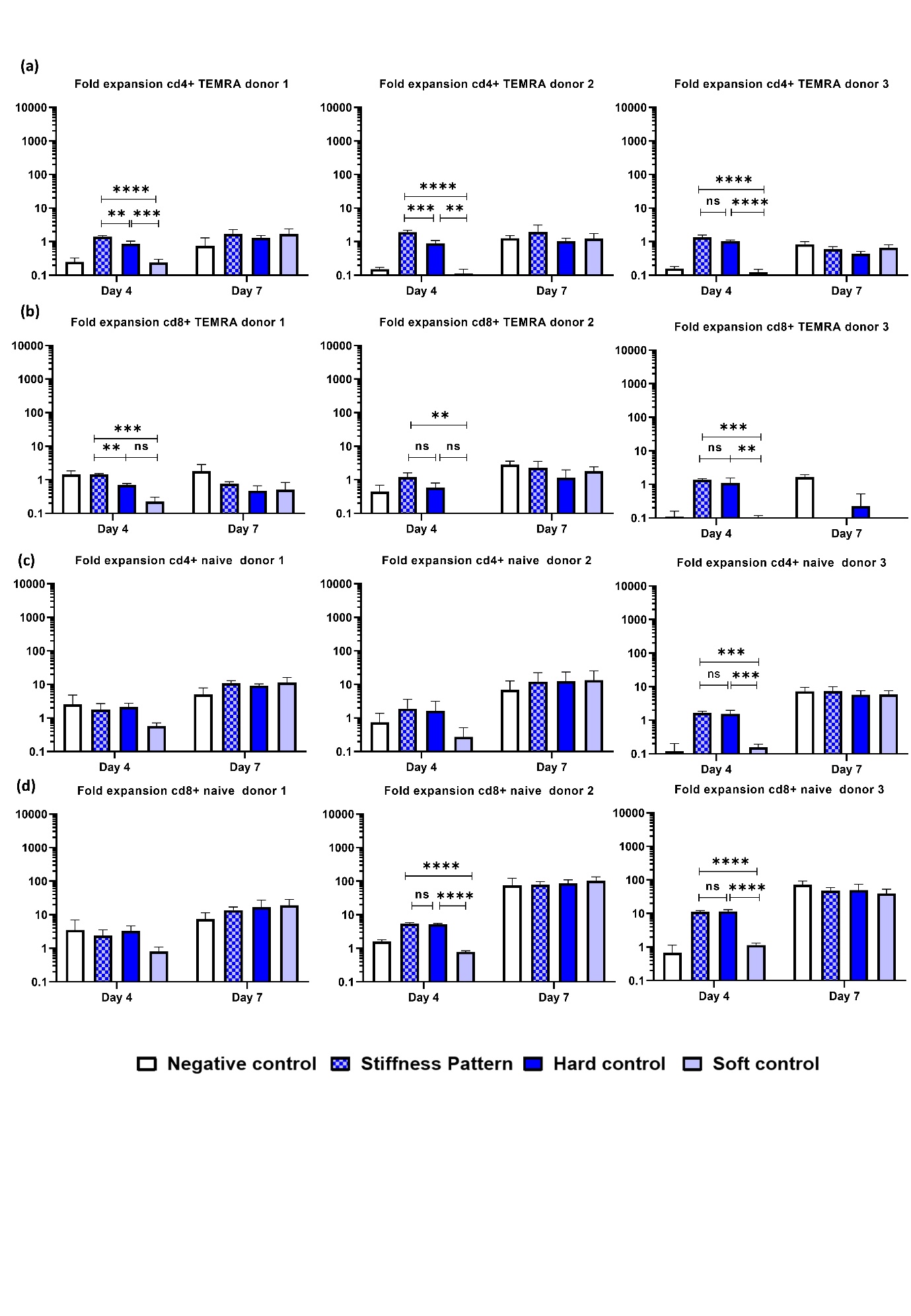

Figure S3. (a) Proliferation of CD4+ TEMRA Phenotype, (b) Proliferation of CD8+ TEMRA Phenotype, (c) Proliferation of CD4+ Naive Phenotype. (d) Proliferation of CD8+ Naive Phenotype. The statistical analysis was performed in Tukey’s multiple comparison test using GraphPad Prism software. *p < 0.05, **p < 0.01, ***p < 0.001, ****p < 0.0001, and ns: not significant .
